# Therapeutic assessment of targeting ASNS combined with L-Asparaginase treatment in solid tumors and investigation of resistance mechanisms

**DOI:** 10.1101/2020.09.08.287565

**Authors:** Verena Apfel, Damien Begue, Valentina Cordo, Laura Holzer, Laetitia Martinuzzi, Alexandra Buhles, Grainne Kerr, Ines Barbosa, Ulrike Naumann, Michelle Piquet, David Ruddy, Andreas Weiss, Stephane Ferretti, Reinaldo Almeida, Debora Bonenfant, Luca Tordella, Giorgio G. Galli

**Affiliations:** Disease area Oncology, Novartis Institute for Biomedical Research, Basel, Switzerland; Analytical Sciences and Imaging, Novartis Institutes for Biomedical Research, Basel, Switzerland; Disease area Oncology, Novartis Institute for Biomedical Research, Cambridge, USA; Global Discovery Chemistry, Novartis Institutes for Biomedical Research, Basel, Switzerland

## Abstract

Asparagine deprivation by L-Asparaginase (L-ASNase) is an effective therapeutic strategy in Acute Lymphoblastic Leukemia, with resistance occurring due to upregulation of ASNS, the only human enzyme synthetizing Asparagine^1^. L-Asparaginase efficacy in solid tumors is limited by dose-related toxicities ^2^. Large-scale loss of function genetic in vitro screens identified ASNS as a cancer dependency in several solid malignancies ^3,4^. Here we evaluate the therapeutic potential of targeting ASNS in melanoma cells. While we confirm in-vitro dependency on ASNS silencing, this is largely dispensable for in vivo tumor growth, even in face of asparagine deprivation, prompting us characterize such resistance mechanism to devise novel therapeutic strategies. Using *ex vivo* quantitative proteome and transcriptome profiling, we characterize the compensatory mechanism elicited by ASNS knockout melanoma cells allowing their survival. Mechanistically, a genome-wide CRISPR screen revealed that such resistance mechanism is elicited by a dual axis: GCN2-ATF4 aimed at restoring amino acids levels and MAPK-BCLXL to promote survival. Importantly, pharmacological inhibition of such nodes synergizes with L-Asparaginase-mediated Asparagine deprivation in ASNS deficient cells suggesting novel potential therapeutic combinations in melanoma.

## Introduction

Asparagine deprivation is a therapeutic strategy long used in clinical setting as part of the chemotherapeutic regimen for Acute Lymphoblastic Leukemia (ALL) by administration of the prokaryotic enzyme L-Asparaginase (L-ASNase) ^1^. While such treatment paradigm is particularly efficacious in ALL, resistance is observed in patients by upregulation of Asparagine Synthetase (ASNS) ^1^. Additionally, dose limiting toxicities hampered the employment of L-ASNase in solid tissues and several alternative enzyme modifications or delivery methods are under continuous investigation ^2^.

Several reports ^5,6^, as well as a large-scale metabolome profiling in various cancer types, highlighted the dependency of cancers displaying low levels of Asparagine (e.g. pancreatic) to ASNS gene silencing ^7^. However, by using large-scale barcoding experiments, we previously reported that Asparagine deprivation affects only a small subset of cell lines bearing *ASNS* promoter hypermethylation ^7^, further challenging the validity of Asparagine deprivation as an impactful strategy for cancer therapy.

ASNS is the only gene encoded in the mammalian genome that converts Aspartate into Asparagine. This is achieved by the ATP-dependent amidation of L-Aspartate using L-glutamine as a nitrogen source. Several reports highlighted a functional role for ASNS in tumor growth and metastatic dissemination in various cancer settings ^5,6,8^, thereby prompting the necessity to identify inhibitors to validate the cancer dependency on the catalytic activity of ASNS and potentially improve the current therapeutic window of Asparagine deprivation treatments by L-Asparaginase administration.

We hereby focus on melanoma as a tumor type in which the potential for ASNS as a therapeutic target is largely unexplored. By generating ASNS knockout (KO) cells, we validate that ASNS depletion results in strong anti-proliferative effects *in vitro*, while being largely dispensable for *in vivo* tumor growth. Proteome- and transcriptome-wide analyses of xenograft tumors identify a complex and concerted compensatory mechanism allowing ASNS KO tumors to survive in face of acute Asparagine deprivation by systemic L-Asparaginase administration. Additionally, by genome-wide CRISPR screens, we identify critical pathway nodes that can be exploited as combinatorial therapeutic strategies with ASNS inhibition and Asparagine deprivation.

## Results

### ASNS KO melanoma cells are sensitive to aminoacid deprivation

ASNS was previously identified as a top hit in large scale functional genomic screens in cells displaying low levels of its enzymatic product, the aminoacid asparagine ^7,9^. Careful evaluation of such datasets reveals that cell lines sensitive to ASNS silencing belong to specific lineages such as Pancreatic, Breast, Colorectal, Sarcomas and Cutaneous Melanomas (Figure S1A). While several reports identified such vulnerability in cancers from pancreas ^10,11^, colon ^7^, soft tissues ^6^ and breast ^5^, to our knowledge the therapeutic potential of ASNS inhibition in melanoma is largely unexplored.

Importantly, also in melanoma, the sensitivity of cancer cell lines to shRNAs against ASNS correlates with lower level of Asparagine as reported for other lineages (Figure S1B).

Here we evaluate the maximal achievable efficacy of ASNS inhibition by generating A2058 ASNS KO cells. We use A2058 as we previously reported that ASNS knockdown in A2058 results in decreased proliferation upon aminoacid deprivation in cell culture (Figure S1B) ^7^ and aminoacid deprivation in such cell line leads to ATF4 dependent regulation of ASNS, suggesting proper pathway regulation (Figure S1C). We used two independent sgRNAs targeting the Asparagine Synthetase domain of ASNS to induce ASNS deletion (Figure 1A). We observed drastic deletion of ASNS in polyclonal cell line but nonetheless we isolated single cell clones to ensure full deletion of ASNS and avoid potential escaper cells outgrowth (Figure 1B).

**Figure 1.**
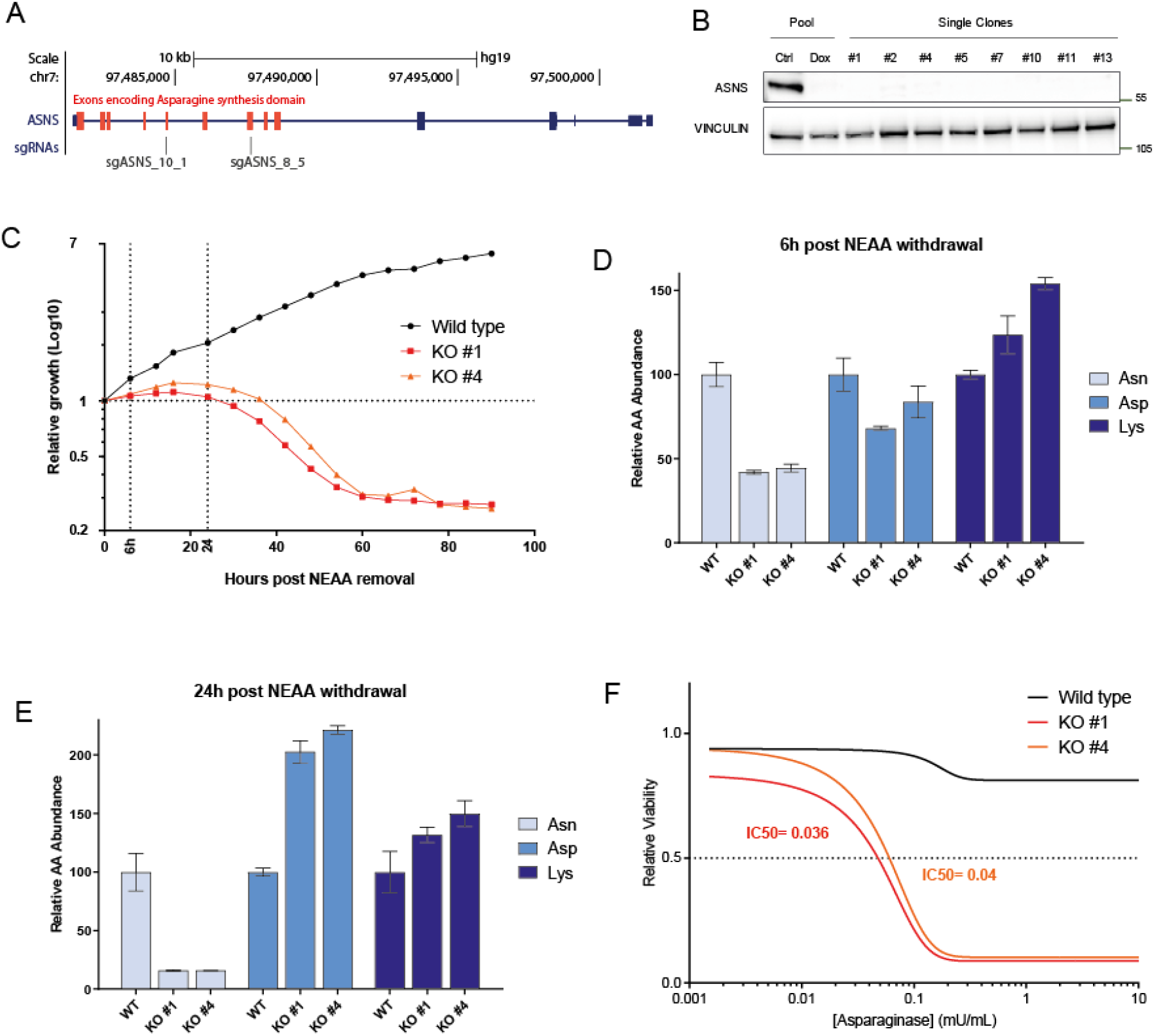
A) UCSC genome browser track encompassing the ASNS locus representing the location of the sequences targeted by the two sgRNAs employed to generate ASNS KO cells. Red boxes represent exons encoding for Asparagine Synthetase domain. B) Western blot analysis of A2058 cells bearing doxycycline inducible sgASNS constructs upon doxycycline treatment (lane 1 and 2). Single cell clones derived from transient expression of sgASNS (lane 3-10). Vinculin is used as a loading control. C) Relative growth (Log10 cell number values as measured by Incucyte – y-axis) for A2058 wild type and ASNS KO clones. X-axis represent the hours elapsed after NEAA deprivation from cell culture media. D-E) Relative abundance of the indicated aminoacids in A2058 WT and KO cells at 6hours (D) and 24 hours (E) removal of NEAA. F) Relative viability (measured by Cell titer Glo – y-axis) of A2058 WT and KO cells upon increasing concentration (x-axis) of L-asparaginase.

ASNS wild type and KO cells were indistinguishable in culture in full media (data not shown), while deprivation of NEAA (Non-essential aminoacids) resulted in decreased proliferation as early as few hours (Figure 1C), with concomitant decrease of cellular Asparagine levels (Figure 1D). Around 24 h post-NEAA deprivation, KO cells started undergoing strong cell death (Figure 1C) matching a further decrease in Asparagine and a concomitant increase in Aspartate (Figure 1E), further corroborating the notion that ASNS is the only enzyme in human catalyzing the conversion of Aspartate to Asparagine.

In order to exclude any potential confounding effect resulting from the depletion of several aminoacids from the NEAA cocktail, we exposed ASNS KO cells to the therapeutic agent L-ASNase. While wild-type cells were largely insensitive to L-ASNase, ASNS KO cells were extremely sensitive in culture (Figure 1F). Our data confirm that ASNS is a key cancer vulnerability in cultured melanoma cells bearing low levels of Asparagine.

### Asparagine depletion does not elicit antitumor response *in vivo*

In order to validate the impact of Asparagine deprivation *in vivo*, we injected A2058 WT and KO cells subcutaneously into mice. We then randomized mice bearing WT and KO cells in two groups in order to treat them with L-ASNase or vehicle control. Importantly, neither in WT, nor in ASNS KO cells, we observed any effect on tumor volume over time compared to control (vehicle) treated group (Figure 2A).Such phenotype (or lack thereof) resembles the lack of response of patients with melanoma and pancreatic cancer to L-ASNase ^12,13^, suggesting the existence of other adaptive survival mechanisms in solid tumors. Tumors assessment by western blot demonstrated that while ASNS WT cells showed the characteristic compensatory upregulation of ASNS upon L-ASNase treatment (Figure 2B), ASNS KO cells, as predicted, did not express any ASNS even upon L-ASNase treatment (Figure 2B).

**Figure 2.**
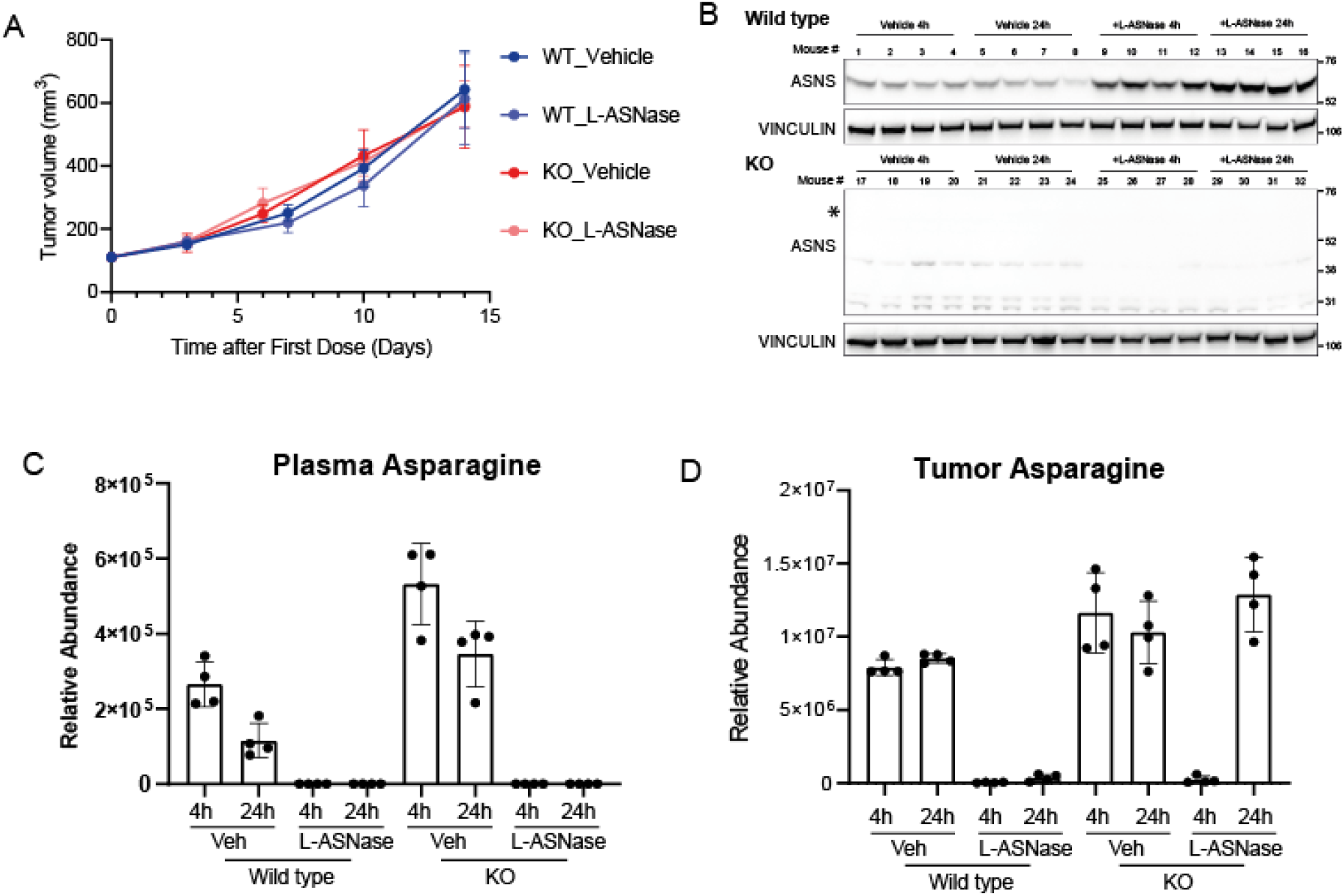
A) Tumor volumes (y-axis) of A2058 WT and KO cells implanted subcutaneously in nude mice and treated or untreated with L-Asparaginase. N= 5 per group and error bars represent SEM. B) Western blot analysis of the tumors harvested from experiment in figure 2A at endpoint. C-D) Relative abundance of Asparaginase in plasma (C) and tumors (D) from mice of experiment in Figure 2A at 4 h and 24 hours after L-asparaginase injection at day 7.

In order to better understand the observed lack of antitumor efficacy, we first evaluated the efficiency of the L-ASNase regimen in depleting circulating asparagine. We then measured Asparagine and Aspartate levels in plasma samples from our experimental cohort by LC-MS 4 h and 24 h post last dosing. In such samples, we observed that L-ASNase treatment reduced Asparagine levels to undetectable levels (Figure 2C) with a concomitant upregulation of Aspartate (Figure S2A). These data suggest that the administration regimen adopted was sufficient to achieve full depletion of circulating Asparagine. We then proceeded to evaluate Asparagine levels in tumors from the same animals. In wild type tumors, L-ASNase treatment significantly suppressed Asparagine levels over the course of 24 h treatment (Figure 2D). Surprisingly, in KO tumors, while Asparagine levels were minimal 4 h post L-ASNase injection, they were restored to the levels of vehicle-treated tumors at 24 hours (Figure 2D). In all samples Aspartate levels were largely unaffected (Figure S2B). Our data suggest that tumors lacking ASNS were endowed with a compensatory mechanism able to re-establish Asparagine levels and sustain growth.

### ASNS KO cells compensatory mechanism consists of deregulation of metabolic and pro-survival signaling pathways

In order to characterize the potential compensatory mechanism elicited by L-ASNase administration in ASNS KO cells, we profiled tumors by quantitative proteomics. To evaluate the specific changes in KO tumors treated with L-ASNase, we compared them with their vehicle treated controls as well as WT tumors treated with L-ASNase. We succeeded to detect and quantitate more than 9000 proteins across the experimental conditions. Interestingly Principal Component Analysis (PCA) analysis revealed that PC1 could distinguish KO-Asparaginase tumors both from WT-Asparaginase and KO-Vehicle tumors (Figure S3A).

Differential expression analysis revealed a large number of proteins either up- or down-regulated in KO tumors treated with Asparaginase (Figure 3A). Gene ontology analysis of upregulated proteins revealed a strong enrichment in categories related to amino acid metabolism (Figure S3B). Indeed we observed significant increase in proteins involved in catalysis of several aminoacids such as PYCR1, PHGDH, KYNU as well as tRNAs such as CARS, WARS etc. and proteins involved in nucleotide metabolism and other metabolic pathways (e.g. NNMT and GMPPA) (Figure S3C).

**Figure 3.**
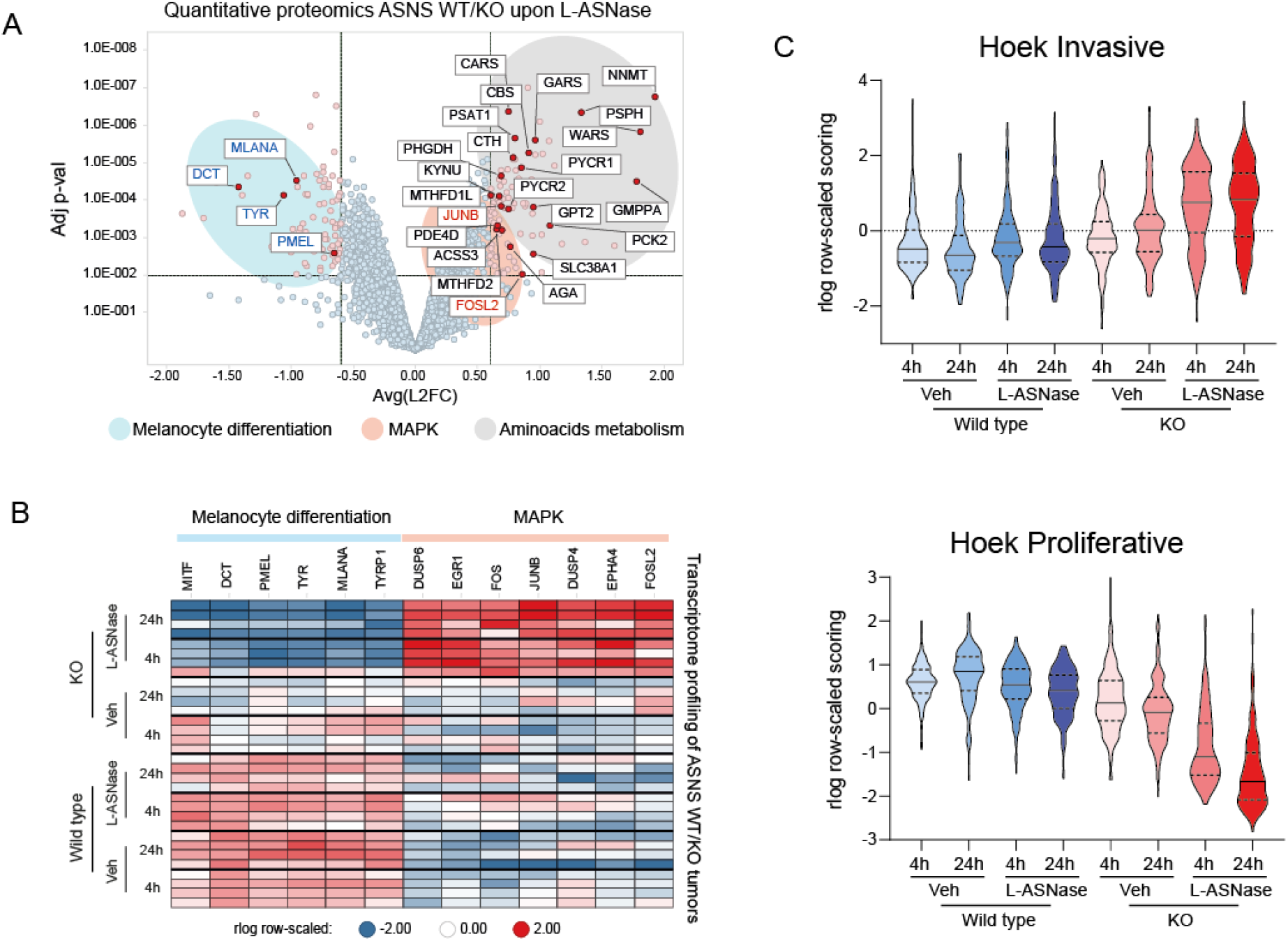
A) Volcano plot representing the Log FC (x-axis) and Adjusted p-value (y-axis) of quantitative proteomics data comparing tumors ASNS KO treated with L-ASNase and matching Wild type tumors with same treatment. (each dot represent the average for each protein. N=4 per group). Black / red / blue gene names represent genes involved in Aminoacid metabolism, MAPK pathway and Melanocyte differentiation respectively. B) Heatmap of RNA expression levels (as measured by RNA-seq) of genes belonging to melanocytic differentiation or MAPK from each individual tumors from experiment in figure 2A. C) Violin plot representing the aggregated expression of signatures reported by ^15^ indicating the “invasive” (top panel) or “proliferative” (bottom panel) status of melanoma tumors.

Interestingly, among deregulated proteins, we observed genes involved in two additional pathways. On one side, several MITF target genes were downregulated, contributing to the gene ontology enrichment for “melanin production”, suggesting cellular de-differentiation (Figure 3A, S3B and S3D). Additionally, among the upregulated proteins, we observed two factors downstream of MAPK signaling such as FOSL2 and JUNB (Figure 3A and S3D).

In order to have a broader analysis of such pathway deregulation, we surveyed the transcriptome of the entire tumors cohort by RNA-sequencing. In agreement with our proteome profiling, we observed several downstream targets of MAPK activity to be upregulated (e.g. EGR1, FOS, DUSP4) while pigmentation genes, prototypically regulated by MITF, were downregulated (e.g. PMEL, TYR) (Figure 3B). These findings suggest a possible transcriptional adaptation to Asparagine deprivation as previously reported for targeted therapies ^14^. To validate these findings, we then applied previously described gene signatures involved in melanoma proliferation and invasiveness ^15^. We observed a concomitant upregulation of the invasiveness signature and downregulation of the proliferative signature in ASNS KO tumors treated with L-Asparaginase (Figure 3C) suggesting that ASNS KO tumors adopt the so-called “phenotype switching” to resist to L-ASNase induced cell death. Our data indicate that upon extracellular amino acid deprivation and in face of the absence of the capacity to synthetize Asparagine, ASNS KO tumors are able to activate multiple pathways as resistance mechanisms for survival.

### GCN2 and MAPK pathways are critical nodes mediating resistance to Asparagine deprivation

In order to identify critical nodes that could be exploited therapeutically to overcome the resistance of ASNS KO cells to Asparagine-deprivation, we performed a genetic loss of function screen in wild type and KO cells upon aminoacid deprivation. We engineered ASNS WT and ASNS KO cells to express Cas9 and performed genome wide CRISPR screens both in full media and in media deprived of Non Essential Amino Acids (NEAA) (Figure 4A).

**Figure 4.**
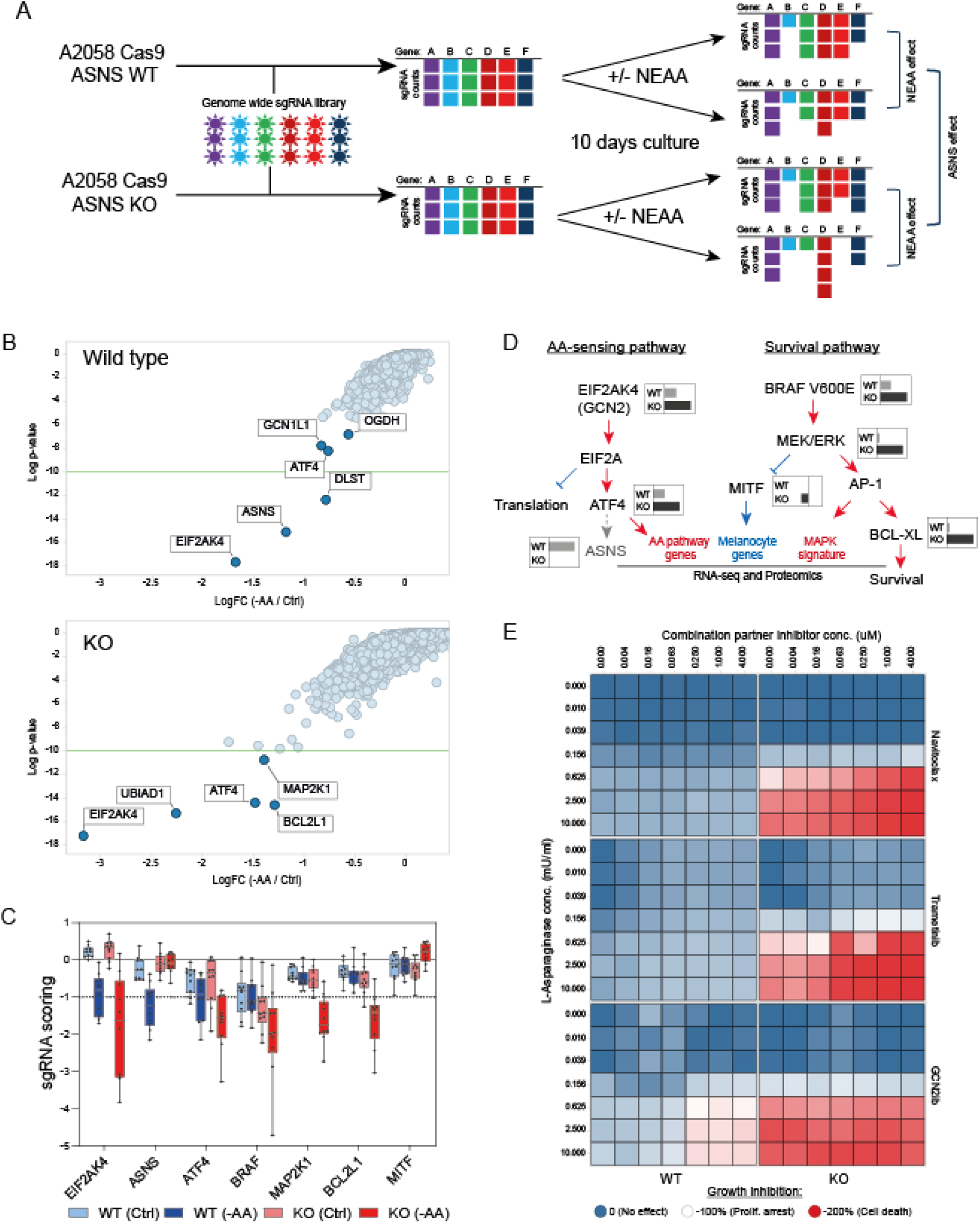
A) Schematic outline of the genome wide CRISPR screen in A2058 ASNS WT and KO cells upon NEAA levels manipulation. B) Results from genome wide CRISPR screens. Data from WT (top panel) and KO (bottom panel) cells were represented as LogFC (x-axis) for each cell line grown either in full media or in NEAA-deprived media. Y-axis represents the Log p-value of significance. C) sgRNA representation (Y-axis) for the individual sgRNAs targeting hits identified in the CRISPR screen from figure 4A and S3E. D) Diagram summarizing findings from CRISPR screening and proteomic and transcriptomic analyses. Key players in AA-sensing and Survival pathways are represented. Grey/black inlets show representative LogFC sensitivity as detected by the CRISPR screen in ASNS WT or KO cells as per figure 4B. Red /blue writings and arrows represent pathways pathways up/down regulated in ASNS KO cells upon L-ASNase treatment. E) Heatmap representing the cell viability (as measured by Cell Titer Glo) of WT (left column) and KO (right columns) A2058 cells treated with increasing concentration of L-Asparaginase (Y-axis) or the indicated compound (X-axis). Colors range between 0 and −200 according to the sensitivity in which −100% corresponds to proliferation arrest.

As expected, wild type cells displayed significant sensitivity to sgRNAs targeting components of the aminoacid sensing machinery upon aminoacid deprivation. Indeed, we observed depletion of sgRNAs targeting the GCN2 kinase (EIF2AK4), its activator GCN1L1 as well as the downstream factors ATF4 and ASNS itself (Figure 4B, top panel). Additionally, we observed sensitivity to two enzymes of the 2-oxoglutarate dehydrogenase complex (OGDH and DLST) (Figure 4B, top panel). In ASNS KO cells, upon NEAA deprivation we observed enhanced sensitivity to targeting EIF2AK4 and ATF4 compared to wild type cells, but also additional scoring for genes such as the prenyltransferase UBIAD1, MAPK effector MAP2K1 (MEK1) and the anti-apoptotic protein BCL2L1 (BCL-XL) (Figure 4B, bottom panel) suggesting rewiring of critical survival pathways in ASNS KO cells. Importantly, these data were supported by multiple independent sgRNAs contained in the library inducing the enhanced sensitivity to the identified hits in ASNS KO cells upon aminoacid deprivation (Figure 4C). The identified hits further corroborate our ex-vivo proteomics and transcriptomics data, with GCN2 and ATF4 controlling aminoacid biosynthesis while MAP2K1 and BCLXL controlling proliferation and survival downstream of MAPK activity (Figure 4D). We were particularly intrigued by the sensitivity of ASNS KO cells to actionable targets such as BCL-XL (inhibited by Navitoclax ^16^) and the kinases MAP2K1 (inhibited by Trametinib ^17^) and GCN2 (inhibited by GCN2ib ^18^). We then performed dose response combination treatments of A2058 wild type and ASNS KO cells with L-Asparaginase and the three compounds. We observed significant an increase in synergy score in ASNS KO cells compared to wild type for Navitoclax (synergy score WT = 0.6, KO = 16.1) and Trametinib (synergy score WT = 3.6, KO = 18.9) (Figure 4E). Interestingly GCN2ib showed already strong synergy with L-ASNase in A2058 wild type cells (synergy score = 11.5) which could not be further enhanced in KO cells (synergy score = 3.2) (Figure 4E) due to the above-mentioned hypersensitivity to L-ASNase (Figure 1F) and in agreement with GCN2 kinase activity being critical in low aminoacid response already elicited by L-ASNase. In summary, genetic and pharmacological inhibition of BCL-XL, MAPK and GCN2 suggest them as candidate combination partners with Asparagine deprivation in vitro.

## Discussion

We here challenge the potential of ASNS as a therapeutic target in melanoma cells as single agent or in combination with acute Asparagine deprivation elicited by L-Asparaginase administration. We demonstrate that, while ASNS KO cells display extreme sensitivity to decreased extracellular aminoacids *in vitro*, they are able to rewire multiple pathways *in vivo* to sustain tumor growth. Such data are in line with recent clinical trials in melanoma showing a lack of efficacy in face of profound Asparagine decreases ^12^.

In our *in vivo* studies we observe no tumor growth inhibition in A2058 cells (reported to have low levels of Asparagine) upon *in vivo* treatment with L-Asparaginase. On the contrary, albeit in different tumor types, others have reported that acute asparagine deprivation *in vivo* is able to elicit antitumor response ^5,10^. One possible explanation could reside in dose scheduling regimen adopted. We use 1000 iU/Kg dose on a 3 days scheduling vs. a daily dosing of 2000 iU/Kg used in these works. Of note, both of these regimens are significantly above the maximum tolerated doses in humans ^19^ and our PD measurements confirm that our regimen is sufficient to suppress Asparagine levels to undetectable levels, as measured by Mass Spectrometry.

Thereby careful considerations of dose scheduling of L-Asparaginase in preclinical models is critical in order to devise potential combination strategies, particularly in the context of the lack of predictability of the levels of Asparaginase suppression necessary to achieve therapeutic efficacy.

An additional consideration regarding preclinical assessment of Asparaginase-based therapies is the constantly evolving scenario of strategies to improve the Therapeutic Index of Asparagine deprivation using Asparaginase from different species (i.e. E. Coli vs. Erwinia Chrysanthemi), modified versions (i.e. Peggylated) or even new delivery methods as for example a cell therapy strategy based on red-blood cells encapsulation ^10^.

Our genome wide CRISPR screens identify therapeutic nodes that might be important to devise novel combination strategies with Asparagine deprivation. During the drafting of this work, the relationship between Asparagine deprivation and MAPK pathway was confirmed by others ^18^ and pressure-tested *in vivo* ^8^. Importantly, in the context of syngeneic mouse models, the authors demonstrate some synergistic effect of inhibiting MAPK and Asparagine, however such effects encompass largely tumor stasis or growth delay rather that potent tumor growth inhibition ^8^. Such data suggest that maybe additional combination partners should be considered. Moreover, the combination between Asparaginase and GCN2 has been mostly explored in vivo in AML models (a liquid tumor indication with high basal level of asparagine) rather than melanoma ^18^.

Our results provides possible explanations for the lack of clinical efficacy of L-ASNase trials in melanoma ^12,13^ and provide a framework for studying combinatorial strategies centered around Asparagine deprivation (using L-Asparaginase or by ASNS depletion) and inhibitors of other signaling pathways.

Indeed, in absence of ASNS (hence the inability to synthetize Asparagine *de novo)* at least 2 adaptive mechanisms come into place: a main one GCN2/ATF4-driven aimed at restoring Asparagine levels, complemented by a MAPK-driven one to inhibit apoptosis via BCL-XL while promoting a de-differentiated state via MITF targets inhibition. Future studies will be needed to understand how to best employ the suggested combination regimens preclinically to maximize effective therapeutic strategies based on Asparagine deprivation in solid tumors.

## Material and Methods

### Cell culture and reagents

A2058 melanoma cells were maintained in DMEM supplemented with 10% Fetal Bovine Serum (Seradigm) and 1X MEM NEAA (Gibco). ASNS KO cells were obtained by lentiviral delivery of Cas9 and two sgRNAs (sequences sgASNS_8 5’-GCAGAGTGGCAGCAACCAAG-3’ and sgASNS_10 5’-GGGATCAGATGAACTTACGC-3’) and isolation of single cell clones by limiting dilution.

Cell growth was measured using Incucyte (Essen Bioscience) and proliferation by Cell Titer Glo (Promega) according to manufacturer’s instructions.

L-Asparaginase was obtained by Cedarlane (CLENZ287) and GCN2 inihibitor by MedChem Express (Cat. HY-112654).

Compound dose response proliferation curves were obtained by treating cells for 72 hours at the indicated doses, unless stated otherwise. The anti-proliferative effect of compound combinations was determined based on a 6×6 matrix and compounds were added using the HP D300 digital dispenser in 384 wells plate. Immunoblotting was performed according to standard protocol. Antibodies used are ASNS (Proteintech 14861), Vinculin (Sigma V9131).

### In vivo xenografts

Animal experiments were approved by the Cantonal Veterinary Office Basel-Stadt and were conducted in accordance with the Federal Animal Protection Act and the Federal Animal Protection Order. Animal experiments were performed at Novartis facilities in adherence to the Association for Assessment and Accreditation of Laboratory Animal Care International guidelines as published in the Guide for the Care and Use of Laboratory Animals, and to Novartis Corporate Animal Welfare policies.

A2058 wild type and ASNS KO clone 4 cells (5 milion cells in HBSS) were subcutaneously injected in the flank of 6-8 weeks old female athymic nude mice (Charles River). When tumors reached a mean tumor volume of around 100-150 mm3 animals were randomized into different treatment groups based on similar tumor size and body weight (n=8/group). L-asparaginase (Cederlane, CLENZ287) was dissolved in 0.9% NaCl and animals were treated with 2 iU/g i.p. 3qw. Tumor size was measured twice a week with a caliper. Tumor volume was calculated using the formula (Length x Width) x π/6 and expressed in mm3. Data is presented as mean ± SEM. At completion of the experiment, tumors were isolated, snap frozen in liquid nitrogen and pulverized for molecular analyses using Covaris CP02.

### Next generation sequencing-based techniques

#### RNA sequencing

RNA was prepared from cells or tumors using RNeasy Mini kit (QIAGEN) and RNA-seq libraries were prepared using TruSeq RNA Library Prep Kit v2 (Illumina) according to manufacturer’s recommendations. Libraries were sequenced on a HiSeq 2500 (Illumina).

#### Genome-wide CRISPR screen

A2058 Wild type and ASNS KO cells were plated in CellSTACK Culture Chambers (VWR) 24 h before infection. On the day of infection, the culture media were replaced with fresh media containing 8 μg ml-1polybrene (Millipore) and a genome wide pooled sgRNA library lentivirus ^20^ at a representation of 1,000 cells per sgRNA with a multiplicity of infection of 0.4. Cells were selected for 3 days in the presence of 2 μg ml-1 puromycin for efficient lentivirus transduction, and an aliquot of cells was collected to validate adequate selection. Cells were then splitted in two with either full media or media deprived of NEAA 0. Cells were further propagated for 10 days to identify sgRNAs enriched or depleted in WT or KO cells upon aminoacid deprivation. An average representation of ≥1,000 cells per sgRNA was maintained at each passage throughout the screen. Following completion of the screen, genomic DNA was extracted by the QIAamp DNA Blood Maxi Kit (Qiagen). DNA sequences containing sgRNA templates were PCR amplified from 100 μg of genomic DNA, and PCR fragments were purified using Agencourt AMpure XP beads (Beckman Coulter Life Sciences). The resulting fragments were sequenced on an Illumina HiSeq 4000 platform.

### Amino Acid analysis by LC-MS

10 μL plasma, 20-30 mg tissue or 5*10E6 cells were mixed with 490 μL methanol/formic acid (98/2, v/v) containing 20 μg/ml caffeine as internal standard, sonicated for 15 min and vigorously mixed at 5°C for 30 min with 2000 rpm on a ThermoMixer (Eppendorf, Hamburg, Germany). Samples were centrifuged for 5 min at 18000 rpm. 200 μL of the liquid phase was transferred into a glass vial for LC-MS analysis.

Chromatographic separation was performed using an Intrada Amino Acid column (150mm * 1mm, 3 μm) (Imtact Corp., Japan) on an Ultimate 3000 LC system (Thermo Fisher Scientific, Bremen, Germany). 5 μl of amino acid extract was injected and separated by binary gradient elution. The mobile phase solvents had the following compositions: acetonitrile/tetrahydrofuran/25mM ammonium acetate/ formic acid (9/75/16/0.3, v/v/v/v), solvent A and acetonitrile/100mM ammonium acetate (20/80, v/v), solvent B. The gradient elution program started with 100% A, maintained for 2 min, increase to 17% B over 2.5 min, further increased to 100% B over 3.5min and kept at 100% B for 3 min. The column was then equilibrated for 2.5 min at 100% A. The separation was performed at 40 °C using a flow rate of 100 μl/min.

The LC system was coupled to a QExactive Plus mass spectrometer (Thermo Fisher Scientific, Bremen, Germany). FTMS spectra were acquired in positive ion mode with a target mass resolution of 70.000 in different narrow mass ranges depending of the amino acid of interest (i.e. Asn, m/z 130 to 136).

### Quantitative Proteomics Analysis

The proteins were extracted from homogenized tissues powders with the PreOmics iST-NHS lysis buffer(#P.O.00026). The samples were then processed using the PreOmics kit following their recommendedprotocol with minor modifications. In brief, the proteins were reduced, alkylated and digested for 2h at 37°C. The peptides were then labelled with TMT reagent (1:4; peptide:TMT label)(Thermo Fisher Scientific).

After quenching, the peptides were purified, and the 11 samples were combined to a 1:1 ratio.

Mixed and labeled peptides were subjected to high-pH reversed-phase HPLC fractionation on an AgilentX-bridge C18 column (3.5 μm particles, 2.1 mm i.d., and 15 cm in length). Using an Agilent 1200 LC system,a 60 min linear gradient from 10 % to 40 % acetonitrile in 10 mM ammonium formate separated the peptidemixture into a total of 96 fractions, which were then consolidated into 24 fractions. The dried 24 fractionswere reconstituted in 0.1 % formic acid for LC-MS3 analysis.

Labelled peptides were loaded onto a 15 cm column packed in-house with ReproSil-Pur 120 C18-AQ 1.9μM (75 μm inner diameter) in an EASY-nLC 1200 system. The peptides were separated using a 120 min gradient from 3 % to 30 % buffer B (80 % acetonitrile in 0.1% formic acid) equilibrated with buffer A (0.1 % formic acid) at a flow rate of 250 nl/min. Eluted TMT peptides were analyzed on an Orbitrap Fusion Lumosmass spectrometer (Thermo Fisher Scientific).

MS1 scans were acquired at resolution 120,000 with 350-1500 m/z scan range, AGC target 2×10^5^, maximum injection time 50 ms. Then, MS2 precursors were isolated using the quadrupole (0.7 m/z window) with AGC 1×10^4^ and maximum injection time 50 ms. Precursors were fragmented by CID at a normalized collision energy (NCE) of 35 % and analyzed in the ion trap. Following MS2, synchronous precursor selection (SPS) MS3 scans were collected by using high energy collision-induced dissociation (HCD) andfragments were analyzed using the Orbitrap (NCE 65 %, AGC target 1×10^5^, maximum injection time 120 ms, resolution 60,000).

Protein identification and quantification were performed using Proteome Discoverer 2.1.0.81 with the SEQUEST algorithm and Uniprot human database (2014-01-31, 21568 protein sequences). Mass tolerance was set at 10 ppm for precursors and at 0.6 Da for fragment. Maximum of 3 missed cleavages were allowed.Methionine oxidation was set as dynamic modification; while TMT tags on peptide N termini/lysine residues and cysteine alkylation (+113.084) were set as static modifications.

The list of identified peptide spectrum matches (PSMs) was filtered to respect a 1% False Discovery Rate (FDR) after excluding PSMs with an average TMT reporter ion signal-to-noise value lower than 10 and a precursor interference level value higher than 50%. Subsequently, protein identifications were inferred from protein specific peptides, i.e. peptides matching multiple protein entries were excluded. A minimum of 2 PSMs per protein was required. The final list of identified proteins was filtered to achieve a 5% FDR. Protein relative quantification was performed using an in-house developed python (v.3.4) notebook. This analysis included multiple steps; adjustment of reporter ion intensities for isotopic impurities according to the manufacturer’s instructions, global data normalization by equalizing the total reporter ion intensity across all channels, summation of reporter ion intensities per protein and channel, calculation of protein abundance log2 fold changes (L2FC) and testing for differential abundance using moderated t-statistics (19) where the resulting p-values reflect the probability of detecting a given L2FC across sample conditions by chance alone. The full list of identified and quantified proteins is included as Supplementary Table 1.

Full dataset is deposited in the PRIDE database with accession PXD015152 ^21^.

**Password:** beUP4FID

### Bioinformatic analyses

#### Quantitative mass spectrometry

Protein identification and quantification were performed using Proteome Discoverer 2.1.0.81 with the SEQUEST algorithm and Uniprot human database (2014-01-31, 21568 protein sequences). Mass tolerance was set at 10 ppm for precursors and at 0.6 Da for fragment. Maximum of 3 missed cleavages were allowed. Methionine oxidation was set as dynamic modification; while TMT tags on peptide N termini/lysine residues and cysteine alkylation (+57.02146) were set as static modifications.

The list of identified peptide spectrum matches (PSMs) was filtered to respect a 1% False Discovery Rate (FDR) after excluding PSMs with an average TMT reporter ion signal-to-noise value higher than 10 and a precursor interference level value lower than 50%. Subsequently, protein identifications were inferred from protein specific peptides, i.e. peptides matching multiple protein entries were excluded. A minimum of 2 PSMs per protein was required. The final list of identified proteins was filtered to achieve a 5% FDR. Protein relative quantification was performed using an in-house developed python (v.3.4) notebook. This analysis included multiple steps; adjustment of reporter ion intensities for isotopic impurities according to the manufacturer’s instructions, global data normalization by equalizing the total reporter ion intensity across all channels, summation of reporter ion intensities per protein and channel, calculation of protein abundance log2 fold changes (L2FC) and testing for differential abundance using moderated t-statistics ^22^ where the resulting p-values reflect the probability of detecting a given L2FC across sample conditions by chance alone. The full list of identified and quantified proteins is included as Supplementary Table 1.

Full dataset is deposited in the PRIDE database with accession PXD015152 ^21^.

**Password:** beUP4FID

#### RNA-sequencing

Sequencing reads were aligned to the human transcriptome [Gencode v25] using Bowtie2 ^23^. Gene-level expression quantities (TPM) were estimated by the Salmon algorithm ^24^. Bioconductor package DESeq2 ^25^was used to analyze these gene expression data: data were log transformed using the rlog function.

#### CRISPR screens

Sequencing reads were aligned to the sgRNA library. For each sample, sgRNA reads were counted. Results from individual samples were scaled for library size and normalized using the TMM method available in the edgeR Bioconductor package ^26^. The log fold change in sgRNA abundance in NEAA deprived versus full media condition was calculated using the general linear model log-likelihood ratio test method in edgeR ^27^. The average log fold change of all sgRNAs targeting each individual gene was defined as the enrichment fold change for the corresponding gene. The significance of the enrichment was assessed using the RSA algorithm ^28^. Briefly, all sgRNAs were ranked according to their log fold change signal. The rank distribution of sgRNAs targeting the same gene was examined, and a *P* value was assigned based on an iterative hypergeometric function. The *P* value indicates the statistical significance of all sgRNAs targeting a single gene being distributed significantly higher in rankings than would be expected by chance.

#### Combo treatment calculations

Compound combination activity was determined based on Loewe dose additivity using a weighted synergy score (SS) calculation (12). As synergy scores do not have a natural scale, they only allow relative comparisons within one experiment where synergy scores exceeding a value of 2 point to meaningful combination activities. Numbers in growth matrices represent the effect of single-agent or dual treatment on cell proliferation relative to DMSO-treated cells (value set as 0) with −100% indicative complete block of proliferation and >100 (max −200) indicative of cell death.

## Acknowledgements

All the authors affiliated with Novartis Institute of Biomedical Research are employees of Novartis. The authors declare no competing interests.

## Supplementary figures

**Figure S1.**
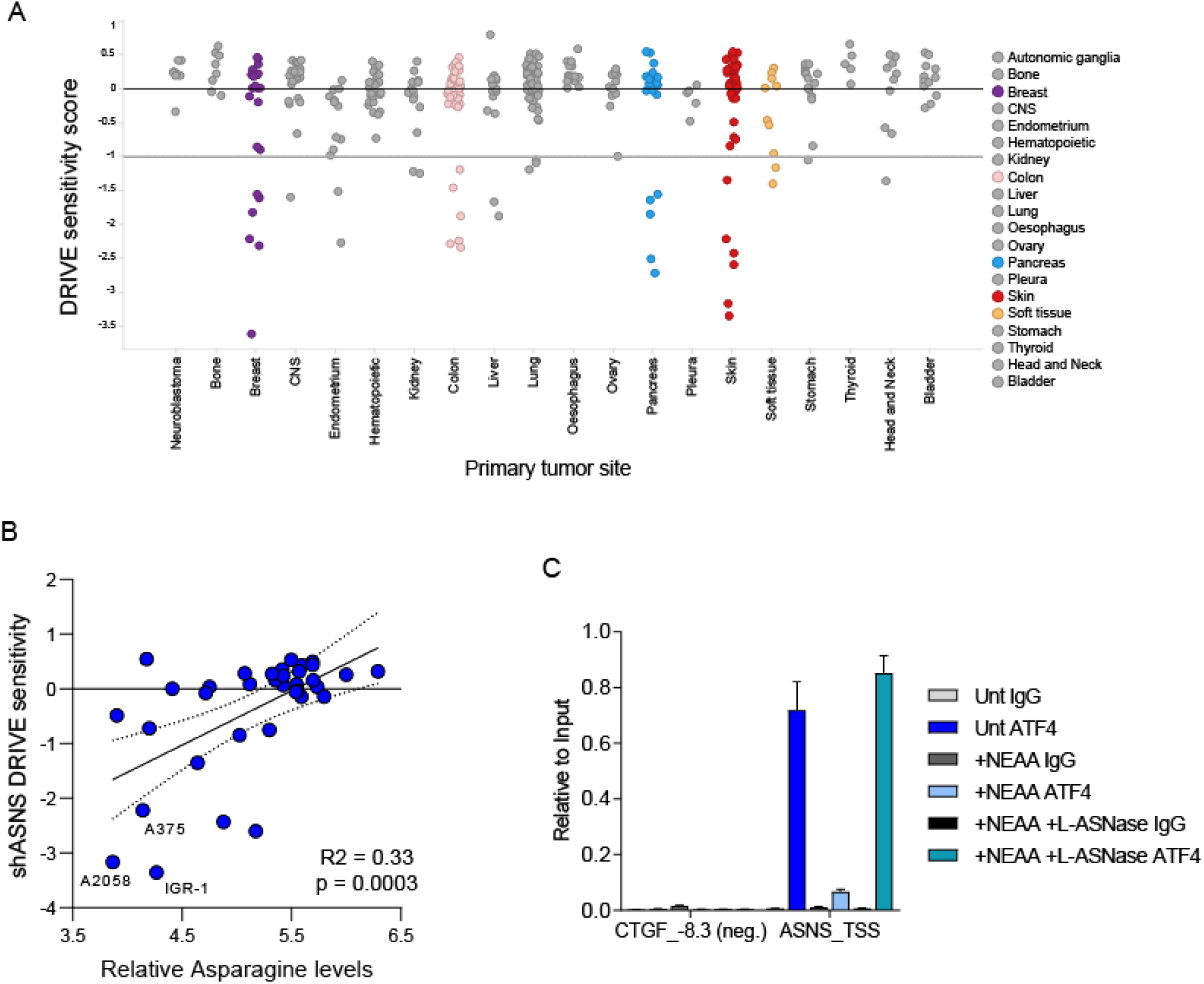
A) Sensitivity score from the DRIVE screen (y-axis) in the panel of cell lines tested binned by cells of origin (x-axis). Horizontal line at −1 represent a treshold of sensitivity as reported in ^9^. B) Correlation between increased sensitivity to shASNS in melanoma cells according to DRIVE dataset and increasing levels of Asparagine as reported in ^7^. C) ATF4 Chromatin immunoprecipitation assay on the ASNS promoter in A2058. Cells were grown either in basic media, supplemented with NEAA or by addition of L-ASNase. CTGF-8.3 is used as a negative control region.

**Figure S2.**
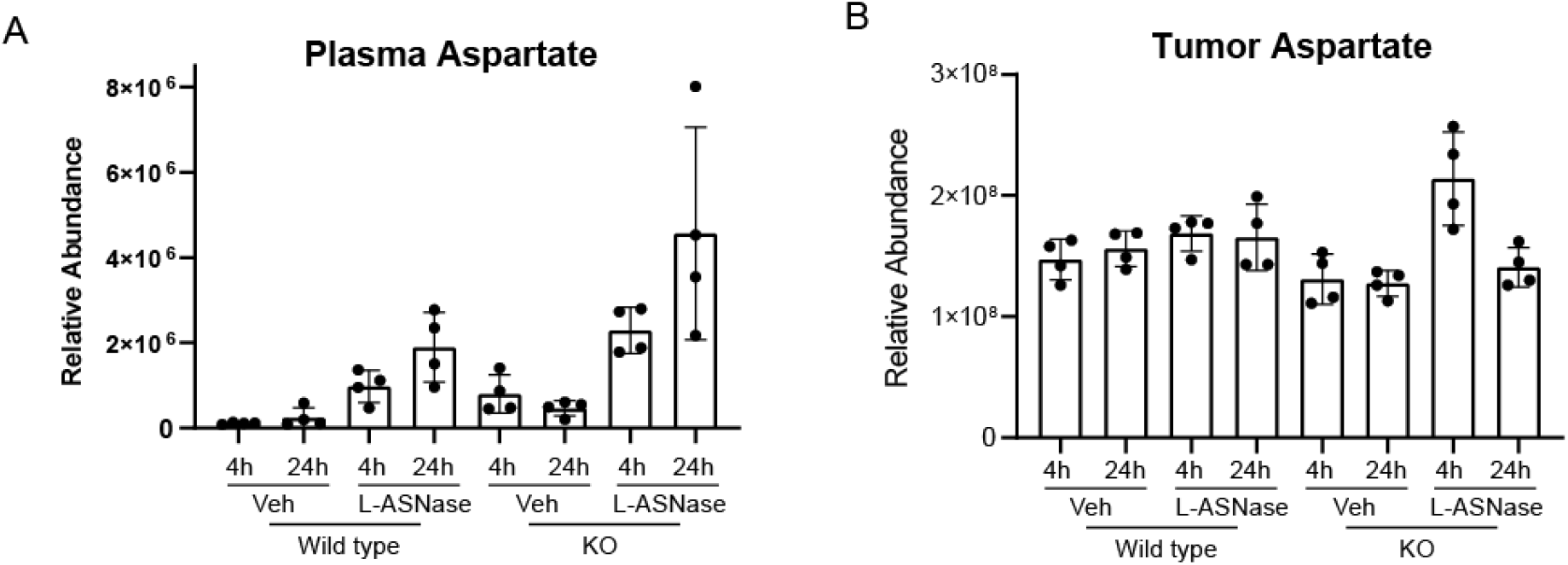
A-B) Relative abundance of Aspartate in plasma (A) and tumors (B) from mice of experiment in Figure 2A at 4 h and 24 hours after L-asparaginase injection at day 7.

**Figure S3.**
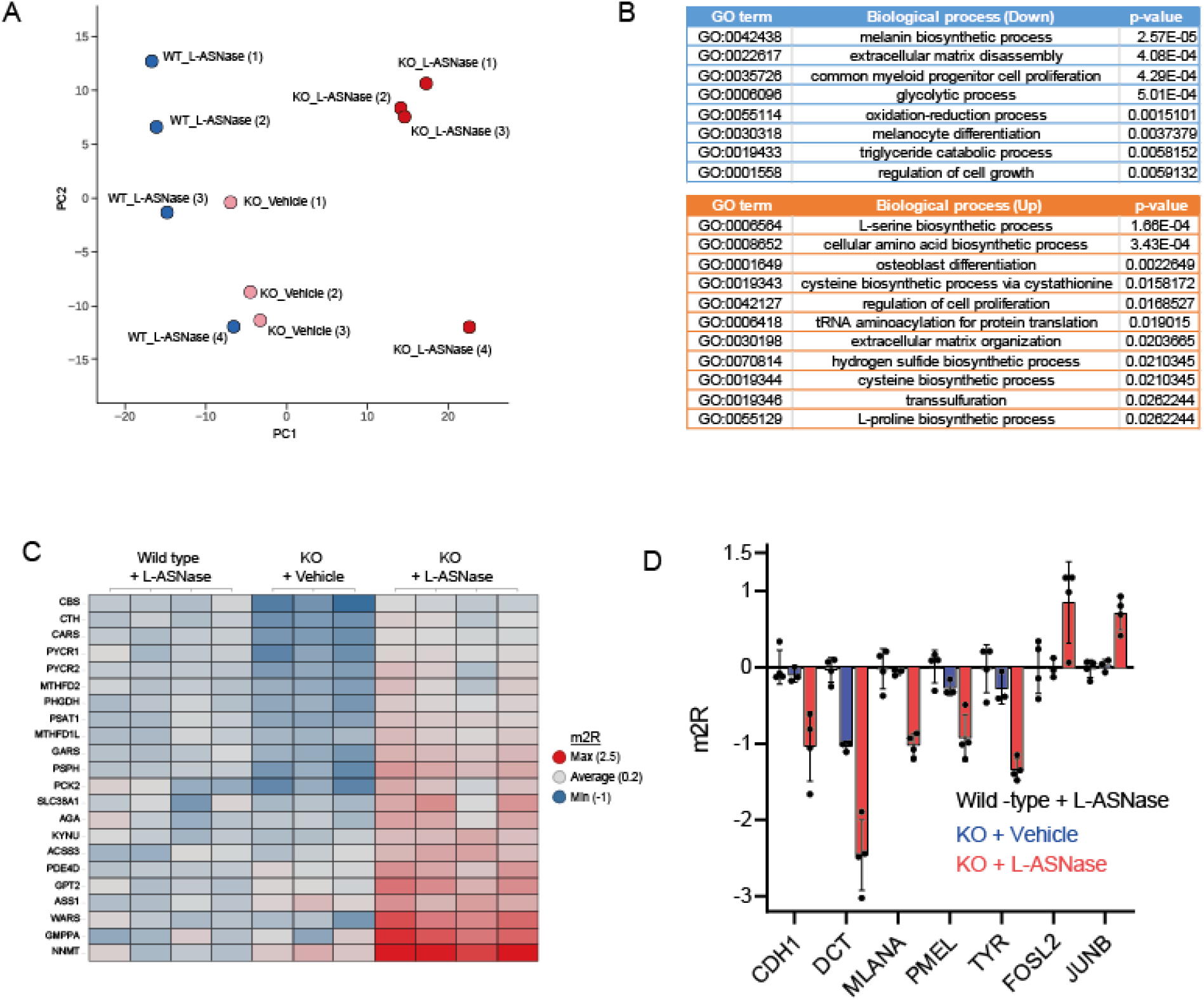
A) PCA plot of the quantitative proteomics data from the analyzed A2058 tumors. Samples are color coded by genotype and treatment. B) Gene Ontology enrichment for proteins differentially expressed in WT and KO tumors treated with L-Asparaginase. C) Heatmap of the proteomics data for differentially expressed proteins involved in metabolic pathways. D) Histogram plot of the proteomics data for proteins involved in melanocyte differentiation (CDH1-TYR) and MAPK (FOSL2 and JUNB).

## References

1. Richards, N. G. J. & Kilberg, M. S. Asparagine Synthetase Chemotherapy. Annu. Rev. Biochem. 75, 629–654 (2006).

2. Koprivnikar, J., McCloskey, J. & Faderl, S. Safety, efficacy, and clinical utility of asparaginase in the treatment of adult patients with acute lymphoblastic leukemia. OncoTargets and Therapy 10, 1413–1422 (2017).

3. Tsherniak, A. et al. Defining a Cancer Dependency Map. Cell 170, 564–576.e16 (2017).

4. McDonald, E. R. et al. Project DRIVE: A Compendium of Cancer Dependencies and Synthetic Lethal Relationships Uncovered by Large-Scale, Deep RNAi Screening. Cell 170, 577–592.e10 (2017).

5. Knott, S. R. V. et al. Asparagine bioavailability governs metastasis in a model of breast cancer. Nature 554, 378–381 (2018).

6. Hettmer, S. et al. Functional genomic screening reveals asparagine dependence as a metabolic vulnerability in sarcoma. Elife 4, (2015).

7. Li, H. et al. The landscape of cancer cell line metabolism. Nat. Med. 25, 850–860 (2019).

8. Pathria, G. et al. Translational reprogramming marks adaptation to asparagine restriction in cancer. Nat. Cell Biol. doi:10.1038/s41556-019-0415-1

9. McDonald, E. R. et al. Project DRIVE: A Compendium of Cancer Dependencies and Synthetic Lethal Relationships Uncovered by Large-Scale, Deep RNAi Screening. Cell 170, 577–592.e10 (2017).

10. Dufour, E. et al. Pancreatic Tumor Sensitivity to Plasma L-Asparagine Starvation. Pancreas 41, 940–948 (2012).

11. Bachet, J.-B. et al. Asparagine Synthetase Expression and Phase I Study With L-Asparaginase Encapsulated in Red Blood Cells in Patients With Pancreatic Adenocarcinoma. Pancreas 44, 1141–1147 (2015).

12. Taylor, C. W., Dorr, R. T., Fanta, P., Hersh, E. M. & Salmon, S. E. A phase I and pharmacodynamic evaluation of polyethylene glycol-conjugated L-asparaginase in patients with advanced solid tumors. Cancer Chemother. Pharmacol. 47, 83–88 (2001).

13. Lessner, H. E., Valenstein, S., Kaplan, R., DeSimone, P. & Yunis, A. Phase II study of L-asparaginase in the treatment of pancreatic carcinoma. Cancer Treat. Rep. 64, 1359–61 (1980).

14. Müller, J. et al. Low MITF/AXL ratio predicts early resistance to multiple targeted drugs in melanoma. Nat. Commun. 5, 5712 (2014).

15. Hoek, K. S. et al. Metastatic potential of melanomas defined by specific gene expression profiles with no BRAF signature. Pigment Cell Res. 19, 290–302 (2006).

16. Tse, C. et al. ABT-263: A Potent and Orally Bioavailable Bcl-2 Family Inhibitor. Cancer Res. 68, 3421–3428 (2008).

17. Gilmartin, A. G. et al. GSK1120212 (JTP-74057) Is an Inhibitor of MEK Activity and Activation with Favorable Pharmacokinetic Properties for Sustained In Vivo Pathway Inhibition. Clin. Cancer Res. 17, 989–1000 (2011).

18. Nakamura, A. et al. Inhibition of GCN2 sensitizes ASNS-low cancer cells to asparaginase by disrupting the amino acid response. Proc. Natl. Acad. Sci. 115, E7776–E7785 (2018).

19. Haskell, C. M. et al. L-Asparaginase. N. Engl. J. Med. 281, 1028–1034 (1969).

20. Liu, H. et al. Tumor-derived IFN triggers chronic pathway agonism and sensitivity to ADAR loss. Nat. Med. 25, 95–102 (2019).

21. Perez-Riverol, Y. et al. The PRIDE database and related tools and resources in 2019: improving support for quantification data. Nucleic Acids Res. 47, D442–D450 (2019).

22. Ritchie, M. E. et al. limma powers differential expression analyses for RNA-sequencing and microarray studies. Nucleic Acids Res. 43, e47–e47 (2015).

23. Langmead, B. & Salzberg, S. L. Fast gapped-read alignment with Bowtie 2. Nat. Methods 9, 357–359 (2012).

24. Patro, R., Duggal, G., Love, M. I., Irizarry, R. A. & Kingsford, C. Salmon provides fast and bias-aware quantification of transcript expression. Nat. Methods 14, 417–419 (2017).

25. Love, M. I., Huber, W. & Anders, S. Moderated estimation of fold change and dispersion for RNA-seq data with DESeq2. Genome Biol. 15, 550 (2014).

26. Robinson, M. D. & Oshlack, A. A scaling normalization method for differential expression analysis of RNA-seq data. Genome Biol. 11, R25 (2010).

27. Robinson, M. D. & Smyth, G. K. Moderated statistical tests for assessing differences in tag abundance. Bioinformatics 23, 2881–2887 (2007).

28. König, R. et al. A probability-based approach for the analysis of large-scale RNAi screens. Nat. Methods 4, 847–849 (2007).

